# System-level studies of a cell-free transcription-translation platform for metabolic engineering

**DOI:** 10.1101/172007

**Authors:** Yong Y. Wu, Hirokazu Sato, Hongjun Huang, Stephanie J. Culler, Julia Khandurina, Harish Nagarajan, Tae Hoon Yang, Stephen Van Dien, Richard M. Murray

## Abstract

Current methods for assembling biosynthetic pathways in microorganisms require a process of repeated trial and error and have long design-build-test cycles. We describe the use of a cell-free transcription-translation (TX-TL) system as a biomolecular breadboard for the rapid engineering of the 1,4-butanediol (BDO) pathway. We demonstrate the reliability of TX-TL as a platform for engineering biological systems by undertaking a careful characterization of its transcription and translation capabilities and provide a detailed analysis of its metabolic output. Using TX-TL to survey the design space of the BDO pathway enables rapid tuning of pathway enzyme expression levels for improved product yield. Leveraging TX-TL to screen enzyme variants for improved catalytic activity accelerates design iterations that can be directly applied to *in vivo* strain development.

## Introduction

Utilizing fast-growing microorganisms to produce molecules of industrial relevance has the potential to rapidly advance the progress of green chemistry. Processes of traditional chemical synthesis require heavy metal catalysts, toxic solvents, and fossil fuels as feedstocks. The biosynthetic approach, which uses naturally occurring enzymes, less energy, and renewable feedstocks, is becoming an attractive alternative.^1–2^ However, biosynthetic approaches are challenged by long design-build-test cycles. Microbial pathway engineering often has about one-week cycle time.^3^ The performance of the pathway is frequently far from design, requiring many iterations.^4^ For example, it took DuPont and Genencor more than 100 person-years of work to develop the commercialization of biobased 1,3-propanediol.^5^ Recent advances in cell-free systems offer an alternative to this costly approach. Cell-free systems have been used to reduce the cycle time of pathway construction. The design-build-test cycle in a cell-free system using linear DNA takes less than one day.^6^

Cell-free systems simulate a controlled cellular environment that delivers repeatable results. Recent research has explored the application of cell-free systems for biological circuits, renewable energy, and medicine. The cell-free transcription-translation (TX-TL) system was first developed as a biomolecular breadboard to test genetic circuits, and many have been demonstrated since.^7–10^ The synthesis of hydrogen and the development of enzymatic fuel cells in cell-free systems has charted new paths for renewable energy.^11–12^ The high yield of therapeutic proteins in *E. coli*-based cell-free synthesis system also offered new methods for medicinal synthesis.^13–15^ Using cell-free systems for prototyping metabolic pathways is an attractive alternative platform for the engineering of biosynthesis in microbial hosts. Lysate of engineered *E. coli* has been used to support high-level conversion of valuable chemicals,^16^ and a cell-free protein synthesis system has been used to screen enzyme variants.^17^ Systematic studies of cell-free systems have monitored the change of pH, measured the change of metabolites over time, and evaluated protein synthesis.^18–20^ Studies have shown that the depletion of ATP is limiting near the beginning of a cell-free reaction, and the consumption of glutamate is critical in regenerating cofactors.^21–23^ A correlation between cell-free and *in vivo* systems has not been demonstrated. As such, a systematic side-by-side analysis between *in vivo* and cell-free systems is required for cell-free systems to be considered widely as a prototyping platform for metabolic engineering.

Metabolic pathways consist of multiple parts and factors that are combined in precise combinations to achieve desired functions. Enzyme expression levels and activity affect target product yield. Tuning enzyme expression requires engineering the level of transcription, translation, and enzyme activity. Furthermore, balancing expression of multiple genes in parallel remains a challenge. Often, enzymes are overexpressed when they are identified to be essential for improving target metabolite production. Protein overexpression can affect cell growth^24^ and possibly reduce metabolite production.^25^ Studies have previously demonstrated the feasibility of tuning protein expression levels *in vivo* for improving metabolite productions.^26–28^ Cell-free TXTL system also provides a platform for investigating the correlation between protein expression levels and metabolite production. Cell-free TX-TL system allows simultaneous protein expression from multiple pieces of DNA, including linear DNA. Such properties can be used to verify pathway enzyme expression and activity. Sun *et al.*’s work has connected protein expression level in TX-TL to *in vivo* systems by comparing the expression strength of different promoters.^6^ This work takes a further step to compare the dynamics of a metabolic pathway in TX-TL to *in vivo* systems by correlating protein expression levels to metabolite production in both systems.

This work aims to demonstrate the reduction of traditional metabolic engineering design-build-test cycles using the 1,4-butanediol (BDO) pathway as a TX-TL prototype. BDO and its derivatives are widely used for producing automotive plastics, electronic chemicals, and elastic fibers. BDO has a projected global market of $8.96 billion by 2019.^29^ Historically, BDO has been produced from petrochemical feedstocks, but recently a bio-based process was commercialized.^30^ The BDO pathway used for this bioprocess is shown in **Figure 1**. The pathway converts a tricarboxylic acid (TCA) cycle intermediate succinyl-coA to BDO. From top to bottom of the pathway schematic, pathway intermediates include succinyl semialdehyde, 4-hydroxybutyrate (4HB), 4-hydroxybutyryl-coA (4HB-coA), and 4-hydroxybutryaldehyde (4HB-aldehyde). CoA-dependent succinate semialdehyde dehydrogenase (*sucD*) catalyzes the conversion of succinyl-coA to succinyl semialdehyde. 4- hydroxybutyrate dehydrogenase (*4-hbd*) catalyzes succinyl-coA to 4HB. 4-hydroxybutyryl-coA transferase (*cat2*) catalyzes 4HB to 4HB-coA. 4-hydroxybutyryl-coA reductase (*ald*) catalyzes 4HB-coA to 4HB-aldehyde. Alcohol dehydrogenase (*adh*) catalyzes 4HB-aldehyde to BDO.

**Figure 1:**
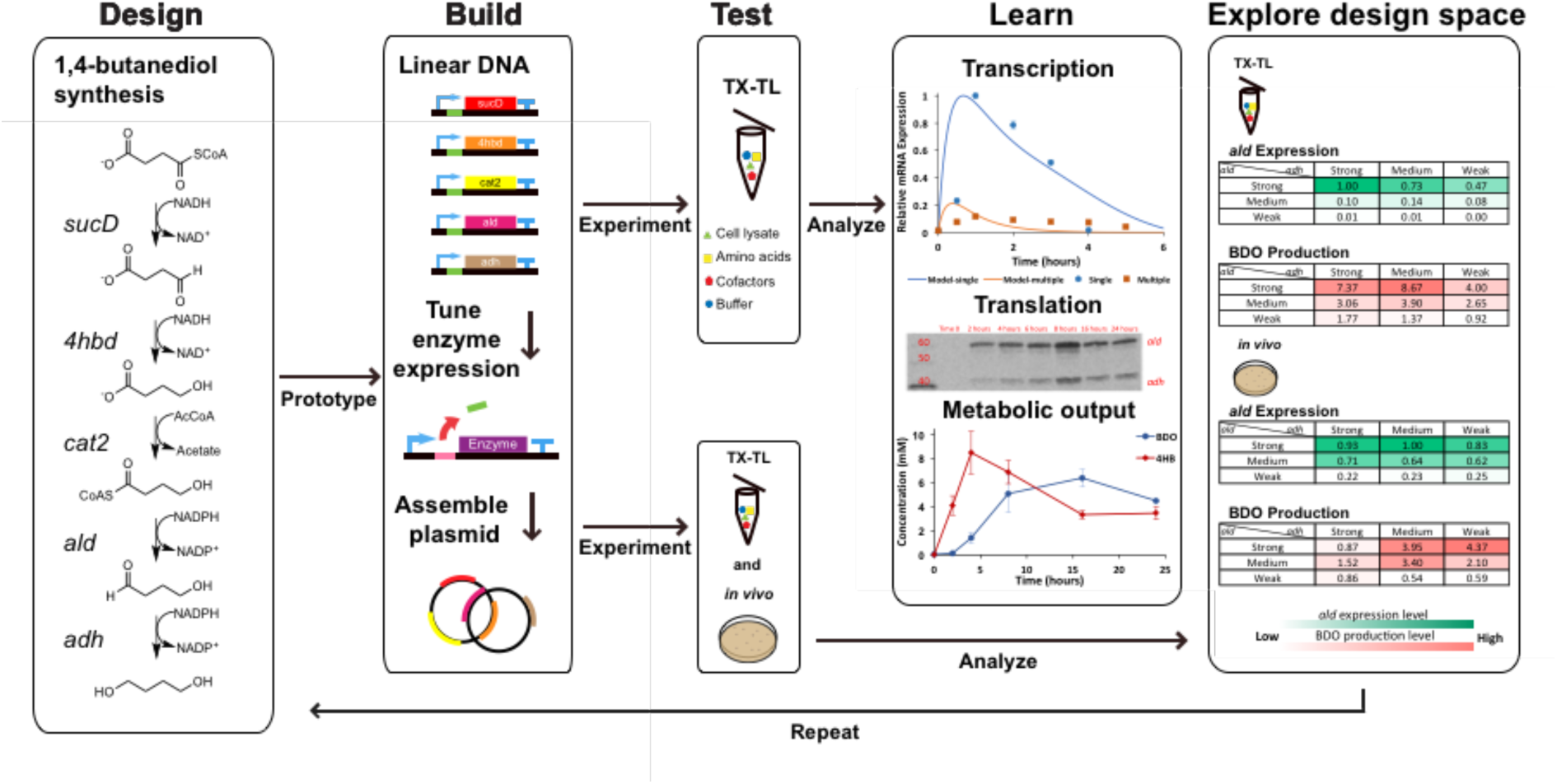
TX-TL as a platform for engineering biosynthetic pathways. A design-build-test-learn cycle using a BDO pathway previously designed by Genomatica Inc. is shown. Linear DNA that encodes pathway enzymes is added to TX-TL. The transcription and translation machinery and metabolic output of TX-TL are analyzed. The design space of the BDO pathway is explored in TX-TL and *in vivo* for comparison. Design: Genomatica Inc. previously designed the BDO synthesis pathway. Build: Generate linear DNA with sequences of pathway enzymes under the same promoters, UTRs, and terminators, subsequently tune enzyme expression levels by varying UTR strength, and assemble into plasmids for design space exploration of the BDO pathway. Test: The TX-TL system is analyzed systematically and compared to the *in vivo* system in parallel. Learn: Systematic analysis of TX-TL through reverse transcription-PCR (RT-PCR), Western blot, and metabolomics. Explore design space: Compare the expression of bottleneck enzyme and BDO production in TX-TL and *in vivo.*

The main goal of this work is to demonstrate TX-TL as a research tool for metabolic engineering and to establish the feasibility of design space exploration (shown in **Figure 1**). Using the BDO pathway previously developed by Genomatica Inc.,^31^ we add linear DNA encoding pathway enzymes to TX-TL. We measure the resulting transcriptional and translational outputs and pathway related metabolites. We also use TX-TL to rapidly tune pathway enzyme expression levels for design space exploration of the BDO pathway. Ribosome-binding site (RBS) elements of varying strengths are chosen from the bicistronic design (BCD) library to adequately explore the design space of the BDO pathway in TX-TL. The two Shine-Dalgarno motifs from BCD allows the first one to make a leader peptide to open the second one, which delivers precise and reliable translation initiation. The translational coupling architecture BCD ensures protein expression at expected levels independent of downstream sequence.^32^ Through exploring the design space of the BDO pathway in TX-TL and *in vivo*, we systematically compare the metabolic output and enzyme expression levels. To show the industrial relevance of TX-TL, we demonstrate that the use of linear DNA in TX-TL has the capabilities to serve as a biomolecular breadboard to speed up design iterations, and results from TX-TL can be applied to *in vivo* strain development.

## Results and Discussion

### Pathway Verification in TX-TL

Pathway enzymes are expressed in TX-TL reactions by adding linear DNA. Linear pieces of DNA are generated by amplifying regions encoding a promoter, a 5’-untranslated region (UTR), a coding sequence of an individual pathway enzyme, and a terminator. End products of TX-TL reactions are directly used for polyacrylamide gel electrophoresis with sodium dodecyl sulfate (SDS-PAGE) preparation, and all enzymes of the BDO pathway show up on the gel at expected sizes (as shown in **Figure S1**). By adding linear DNA encoding enzymes from the BDO pathway *sucD* (*035*), *4hbd* (*036*), *cat2* (*034*), *ald* (*025B*), and *adh* (*012*), we identify that the conversion from 4HB-coA to downstream metabolites is limiting the production of BDO. After 16-hour reactions, more than 10 mM of 4HB is detected, and only 0.5 (± 0.1) mM of BDO is detected. 3.1 (± 0.1) mM of *gamma*-butyrolactone (GBL) is also detected. GBL is the lactonized form of 4HB, and it is hypothesized to be produced spontaneously from 4HB-coA via *cat2*. The production of 4HB and GBL suggests that *sucD* (*035*), *4hbd* (*036*), and *cat2* (*034*) are not rate-limiting enzymes for the pathway, and *ald* (*025B*) and *adh (012*) can be rate-limiting. Previous results from Yim *et al.*^33^ and Barton *et al.*^34^ also agree with our observation of such pathway dynamics. We, therefore, hypothesize *ald* and *adh* as the bottleneck enzymes for the production of BDO in TX-TL. To better understand TX-TL as a platform for metabolic engineering, we pick a more advanced *cat2, ald* and *adh* combination, cat2 (*C*), *ald* (*C*), and *adh* (*C*), for studying a wider range of system dynamics. During an initial experiment with the advanced enzymes, 3.4 (± 0.9) mM of BDO is detected, and about 3.2 (± 0.5) mM of GBL is detected.

The conversion from 4HB to BDO requires the TX-TL system to supplement for electron transfer via cofactors. We subsequently test cofactor concentrations for directing metabolic flux into the production of BDO. We focus on cofactors directly related to the last three steps of the BDO pathway: NADP, NADPH, acetyl-coA, and coA. The absence of acetyl-coA or coA added into the TX-TL system results in less BDO, less GBL, but more 4HB. Also, the addition of acetyl-coA instead of coA helps produce more GBL, which translates to more 4HB-coA synthesized. We hypothesize that the TX-TL system needs more acetyl-coA for the BDO pathway. 1 mM of acetyl-coA is added for the rest of the work. The absence of NADP or NADPH does not drastically affect target metabolite production, but the addition of NADPH improves the production of BDO. 4 mM of NADPH is added for rest of the work, and details can be found in **Figure S2.** These results suggest that the availability of NADPH in TX-TL is limiting the synthesis of downstream products. Such indicates that more strain engineering around the TCA cycle or the pentose phosphate pathway (PPP) may help resolve cofactor imbalance in TX-TL.

### System-level studies in TX-TL

Results from system-level studies of TX-TL system are shown in **Figure 2**. For the analysis of transcription and translation, we focus on the bottleneck enzyme *ald* (*C*). The characterization of the transcription and translation of *ald* (*C*) are carried out by adding linear DNA encoding individual pathway enzymes. We also perform metabolomics on TX-TL reactions to grasp an understanding of the metabolic network.

**Figure 2:**
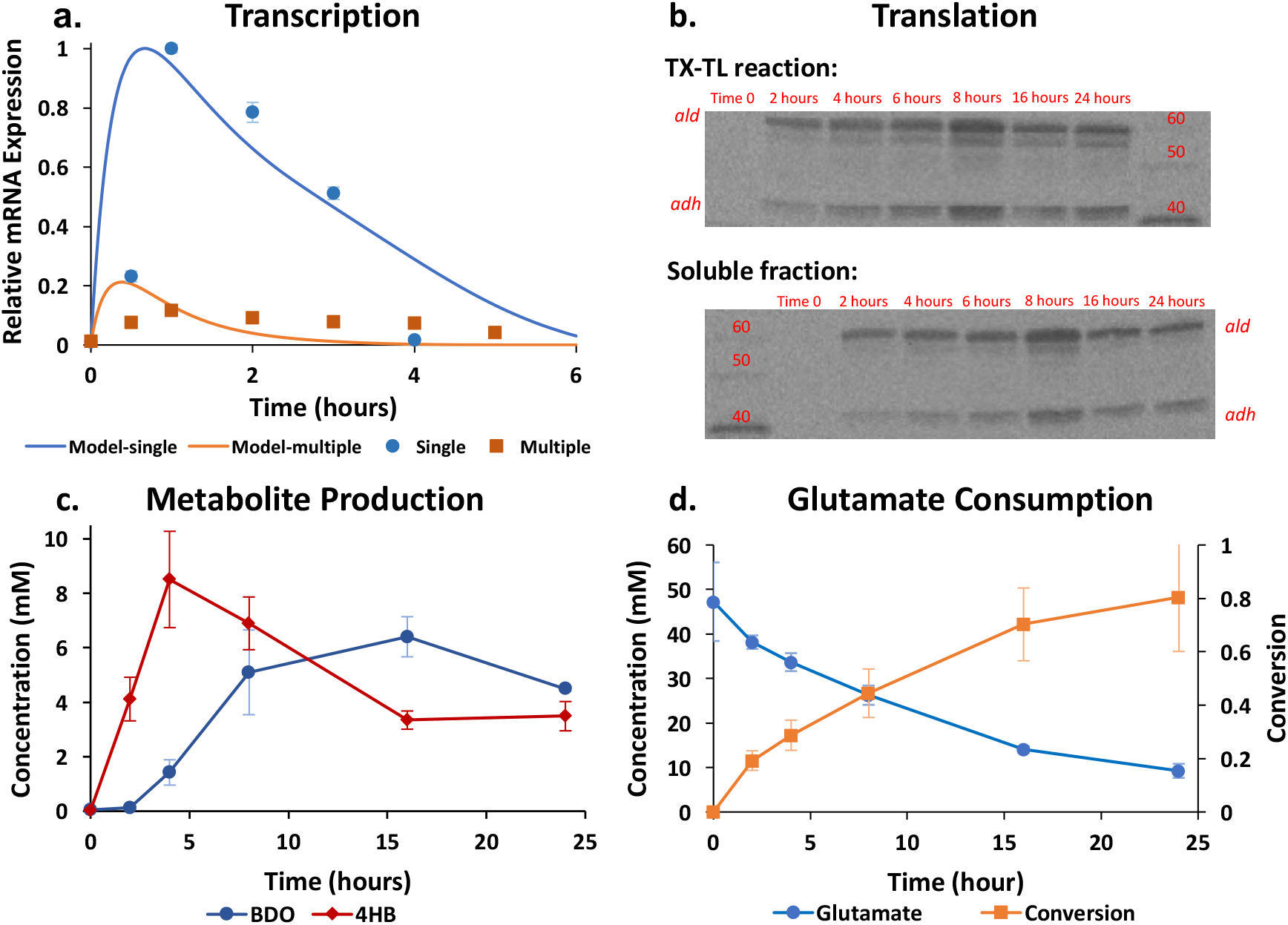
System-level studies of TX-TL. a. Transcription: Relative mRNA for *ald* (*C*) measured using RT-PCR in the first five hours of TX-TL reactions. Blue circles represent data points for mRNA expression of *ald* (*C*) from TX-TL reaction added with just linear DNA encoding *ald* (*C*), and orange squares represent data points for mRNA expression of *ald* (*C*) from TX-TL reaction added with linear DNA encoding all individual pathway enzymes. The blue and orange line represents the respective predictive dynamics. b. Translation: Protein expression analyzed using Western blotting over a 24-hour time range. The top half shows results from analysis of TX-TL reactions, and the bottom half shows results from analysis of supernatant of TX-TL reactions. Both were generated from the same blot. c. Metabolite Production: Measured concentrations of 1,4-BDO (blue circle) and 4HB (red diamond) over time in TX-TL. d. Glutamate Consumption: Glutamate consumption over time in TX-TL (blue circle) and conversion (orange square). Error bars show one standard deviation for n ≥ 3 independent experiments.

#### Transcription

We predict and observe resource limitations of TX-TL by studying the change in mRNA expression dynamics based on how much DNA is added to the TX-TL reaction. **Figure 2a** shows the mRNA expression of *ald* (*C*) from TX-TL reaction added with just linear DNA encoding *ald* (*C*) in blue circles. The orange squares show expression of *ald* (*C*) from TX-TL reaction added with linear DNA encoding individual pathway enzymes. The blue and orange line are the respective predicted mRNA dynamics generated by the TX-TL modeling toolbox.^35^ The linear DNA encoding other pathway enzymes competes with linear DNA encoding *ald* (*C*) for transcription. The mRNA expression difference of the peak mRNA value is greater than 80%. Note that 30 nM of linear DNA encoding *ald* (*C*) is added, and a total of 60 nM of linear DNA encoding other pathway enzymes is added. The transcription of *ald* (*C*) is analyzed using RTPCR. The background signal from DNA is subtracted. The mRNA level peaks within the first hour and then drops to zero by the end of the first five hours. The rapid degradation of mRNA in TX-TL reflects resource limitations of the system. The dynamics of mRNA in TX-TL is similar to previously reported.^36^

#### Translation

The expression of *ald* (*C*) and *adh* (*C*) in TX-TL is analyzed using Western blots and can be normalized by total protein intensities measured from SDS-PAGE.^37^ Complete SDS-PAGE and Western blots for the time-course *ald* (*C*) expression is shown in **Figure S4**. Protein degradation is observed, which is most likely due to the presence of protease in the extract. Extract used in this work is prepared with S138 (MG 1655 Δ*adhE* Δ*ldhA* Δ*pflB* + *lpdA*\*), an engineered *E. coli* strain previously reported.^33^ Most cell-free systems are either developed with purified reagents or with cell lysates from strains with protease deletion. Protein degradation is rarely captured. However, protein degradation is expected here because of the lack of protease gene knockout. The total and soluble protein expression of *ald* (*C*) and *adh* (*C*) in TX-TL are shown in **Figure 2b**. Their respective sizes are 60 kDa and 40 kDa. Most of the enzymes remain in the soluble fraction of the TX-TL reactions. The amount of each protein band on the ladder is approximately 200 ng,^38^ and the concentration of *ald* (*C*) should be on the order of 10 ng/ul.

#### Metabolomics

To understand the metabolism in TX-TL, we carry out experiments to collect time-course data of metabolites. The production of pathway intermediate 4HB and target metabolite BDO is shown in **Figure 2c**. The production of 4HB peaks around 4 hours into the reaction. The level of 4HB drops as the compound is converted to downstream metabolites. The concentration of BDO increases as the concentration of 4HB decreases. The concentration of 4HB-coA peaks between 6 and 8 hours, when BDO concentration starts to plateau, and byproducts GBL starts to accumulate (shown in **Figure S3d**). We hypothesize that the consumption of glutamate links to cofactor regeneration. The production of BDO depends on two steps of NADPH-dependent electron transfer. The concentration of NADPH drops to the detection limit by the end of the fourth hour into the TX-TL reaction. Since NADPH is a critical cofactor for the synthesis of BDO, the rate of NADPH being reduced should be the same magnitude as the rate of NADPH being oxidized. The ratio of NADPH/NADP is shown in **Figure S3b**. Carbon flux through the TCA cycle is very weak, but the consumption of glutamate is very prominent. In **Figure 2d**, the conversion of glutamate reaches 80% by the end of TX-TL reactions. A preliminary ^13^C analysis also confirms that glutamate consumption is the primary energy source (data not shown). Furthermore, the conversion from glutamate to *α*-ketoglutarate reduces NADP to NADPH. The amount of glutamate consumed equals to roughly the sum of 4HB produced, GBL produced, and the BDO produced. The consumption of glutamate is the primary metabolic flux in the TX-TL system.

We learn from the system-level studies that the first few hours are valuable for comparison between pathways or enzyme variants. TX-TL is a resource-limited system: mRNA degradation starts after the first hour, and protein degradation and cofactor imbalance happens. Metabolic flux in TX-TL mainly comes from glutamate, which is also the energy source for regenerating cofactors.

### Design space exploration

The expression levels of *ald* (*C*) and *adh* (*C*) are modulated to explore the design space of the BDO pathway. The expression levels of *sucD* (*035*), *4hbd* (*036*), and *cat2* (*034*) are held at fixed levels. The expression levels of all pathway enzymes are adjusted by tuning the UTR. Constructing the BDO pathway using parts from the BCD library helps modulate pathway enzyme expression levels in TX-TL and *in vivo*. Details of construct design can be found in **Figure S5**. We generate constructs with varying enzyme expression levels, and the level of soluble *ald* (*C*) expression in TX-TL and *in vivo* are shown in **Figure 3a** and **Figure 3d**, respectively. The SDS-PAGE and Western blot images are shown in **Figure S6**. BCD 2, BCD 20, and BCD 22 are used to modify protein expression level to high, medium, and low. We have constructed plasmids with *ald* (*C*) and *adh* (*C*) using a convergent orientation to minimize genetic context effects.^39^ The nine different constructs (shown in **Table S1**) show a range of protein expression levels in TX-TL and *in vivo*. The *ald* (*C*) expression levels are similar in TXTL and *in vivo*. However, the expression of *ald* (*C*) with constructs containing BCD 22 show higher relative expression *in vivo* versus in TX-TL.

**Figure 3:**
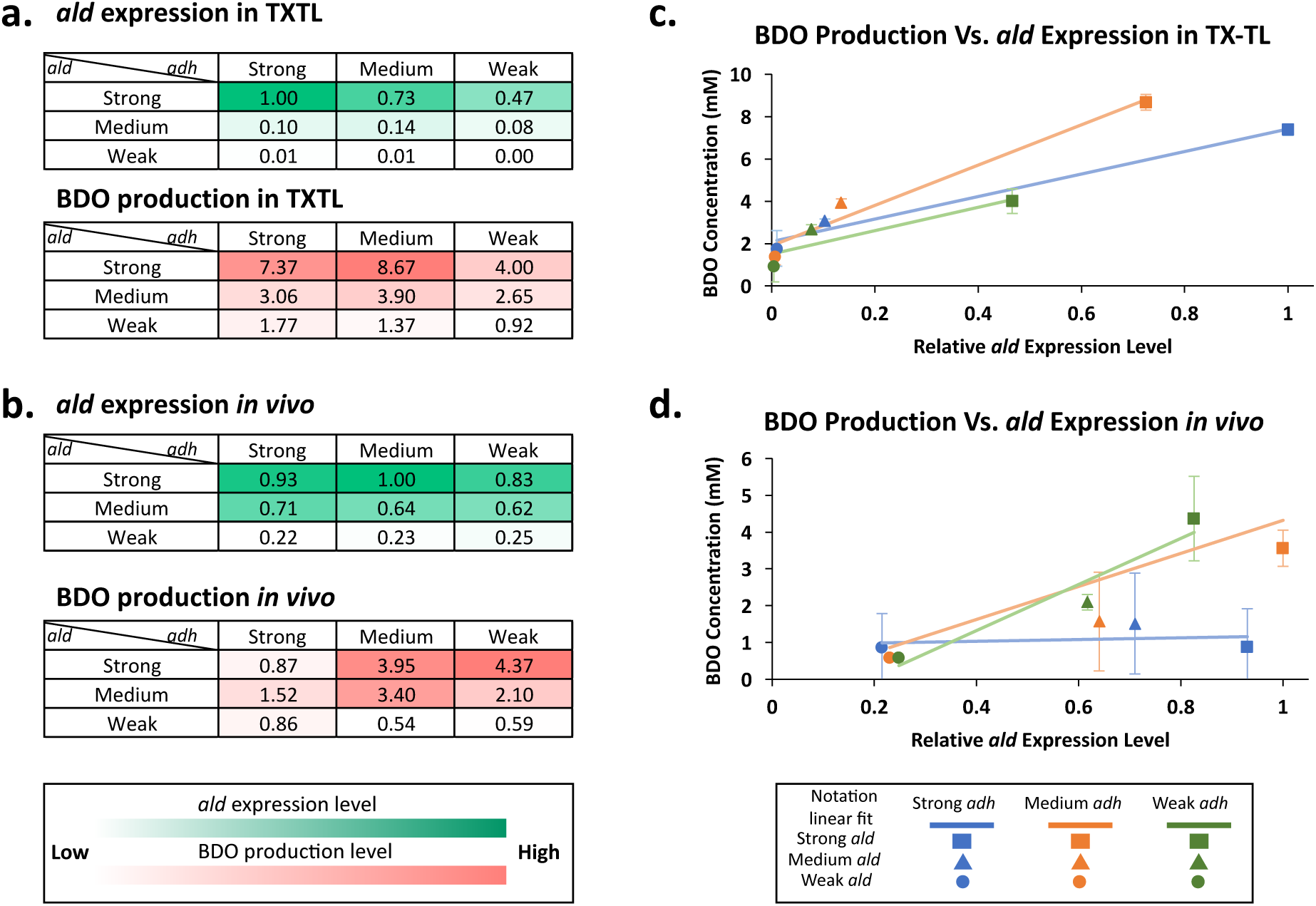
Design Space Exploration in TX-TL and *in vivo.* a. Heat map of *ald* (*C*) expression in TX-TL: Green—*ald* (*C*) expression as the RBS strength of the *ald* (*C*) varies. b. Heat map of the corresponding BDO production in TX-TL: Orange—BDO production (mM) as the RBS strength of the *ald* (*C*) varies. c. BDO production versus protein expression in TX-TL: blue—strong BCD for *adh* (*C*), orange—medium BCD for *adh* (*C*), green—weak BCD for *adh* (*C*), circle: weak BCD for *ald* (*C*), triangle: medium BCD for *ald* (*C*), square: strong BCD for *ald* (*C*). d. Heat map of *ald* (*C*) expression *in vivo*: Green—*ald* (*C*) expression as the RBS strength of the *ald* (*C*) varies. e. Heat map of the corresponding BDO production *in vivo*: Orange--BDO production (mM) as the RBS strength of the *ald* (*C*) varies. f. BDO production versus protein expression *in*

We examine the effect of the expression level of *ald* (*C*) on metabolite production in both systems. Although the metabolic output in TX-TL is not exactly the same as the one *in vivo*, we observe that the production of BDO is closely related to the expression level of the bottleneck enzyme *ald* (*C*) in TX-TL and *in vivo,* which matches with previous studies.^25^ The BDO production level from the constructs in the two systems is different. **Figure 3b** and **3e** shows the production levels of BDO from the designed constructs. The BDO production from constructs with strong *adh* (*C*) expression are lower *in vivo* than in TX-TL. As shown in **Figure 3b** and **3c**, BDO production level is linearly correlated to *ald* (*C*) expression from constructs containing the same BCD in front of the *adh* (*C*) coding region. Overall, the concentration of BDO produced is linearly correlated to the relative expression levels of *ald* (*C*) in TX-TL regardless of the expression level of *adh* (*C*). As shown in **Figure 3e** and **3f**, the concentration of BDO produced is linearly correlated to the relative expression levels of *ald* (*C*) *in vivo* for some constructs. The expression of *adh* (*C*) is overexpressed for constructs containing BCD 2 in front of the coding region of *adh* (*C*). Protein overexpression seems to cause a metabolic burden so strong with high *adh* (*C*) expression that the production of BDO is limited to about 1 mM regardless of the expression level of *ald* (*C*). Data collected from both systems indicates that *ald* (*C*) is the bottleneck enzyme of the BDO pathway.

Resource limitation is a more prominent problem *in vivo* versus TX-TL. The *in vivo* system carries a larger metabolic burden with more complicating factors such as cell growth and the development of antibiotic resistance. While developing and maintaining the transcription and translation machinery, the *in vivo* system also fights against the antibiotics in the culture broth. Although we have used promoter pA^40^ as part of the constructs, leaky enzyme expression is still a problem. Transformation of constructs containing BCD2 (for strong expression) is problematic. At 37 °C, the production of BDO *in vivo* is much lower due to metabolic burden (data not shown). At 30 °C, the production of BDO *in vivo* is comparable to the production of BDO in TX-TL. The production level difference can be the supplement of NADPH and acetyl-coA in the TX-TL system.

The mapping of design space of metabolic pathway from TX-TL to *in vivo* can be further tuned by repeating the design-build-test cycle shown in **Figure 1**. There are two ways to map the two systems more closely. The first way is to redesign plasmid constructs. Re-constructing the pathway with a different set of BCDs, perhaps using BCD 9 or BCD12 instead of BCD 2, can alleviate the metabolic burden *in vivo*. Using weaker BCDs can potentially avoid overexpression enzymes *in vivo,* which can more effectively show the linear correlation between *ald* (*C*) expression level and BDO production. The second way is to tune protein expression in TX-TL. Plasmid concentration within cells are hard to control, but **t**he expression level of proteins in TXTL can be tuned by adjusting the concentration of added DNA.^6^ In other words, the concentration of the added DNA can be adjusted to map protein expression levels in TX-TL to *in vivo*. In this work, we have chosen a relatively low DNA concentration to show a linear range of dynamics between *ald* (*C*) expression and BDO production. More design iterations will lead to better mapping results.

This work has only used batch mode reaction to compare the two systems strictly. However, both systems can benefit from pH control by feeding base. Cell growth is limited by pH (shown in **Figure S9**), and the pH can be tuned by adjusting buffer.

*vivo:* blue—strong BCD for *adh* (*C*), orange—medium BCD for *adh* (*C*), green—weak BCD for *adh* (*C*), circle: weak BCD for *ald* (*C*), triangle: medium BCD for *ald* (*C*), square: strong BCD for *ald* (*C*). Error bars show one standard deviation for n ≥ 3 independent experiments.

### Application of TX-TL

We apply the TX-TL system to compare three different enzyme combinations of the BDO pathway. The reaction conditions for testing the enzyme combinations are the same. The same *sucD* (*035*) and *4hbd* (*036*) are added into TX-TL. The three different enzyme combinations are A: *cat2* (*034*), *ald* (*025B*), and *adh* (*012*) previously published,^33^ B: *cat2* (*B*), *ald* (*B*), and *adh* (*B*) previously published,^34^ C: evolved version of combination B, *cat2* (*C*), *ald* (*C*), *adh* (*C*). As shown in **Figure 4**, we study the production of BDO, 4HB, and GBL in TX-TL. Combination C results with the greatest BDO production rate, and the production of byproduct GBL is the lowest respectively. The production of 4-HB is very similar for all three combinations, but this could be the conversion to 4-HB is reaching equilibrium by the end of the TX-TL reaction. Since the conversion from glutamate to 4-HB is redox neutral, the accumulated 4-HB is almost the same for all three combinations. Enzyme expression comparison between Combination B and Combination C is shown in **Figure 4b**. Notably, the expression of *ald* and *adh* in combination B is much weaker than the expression of *ald* and *adh* in combination C. The engineered enzymes from Combination C is evolved for stability and enzyme specificity for NADPH.

**Figure 4:**
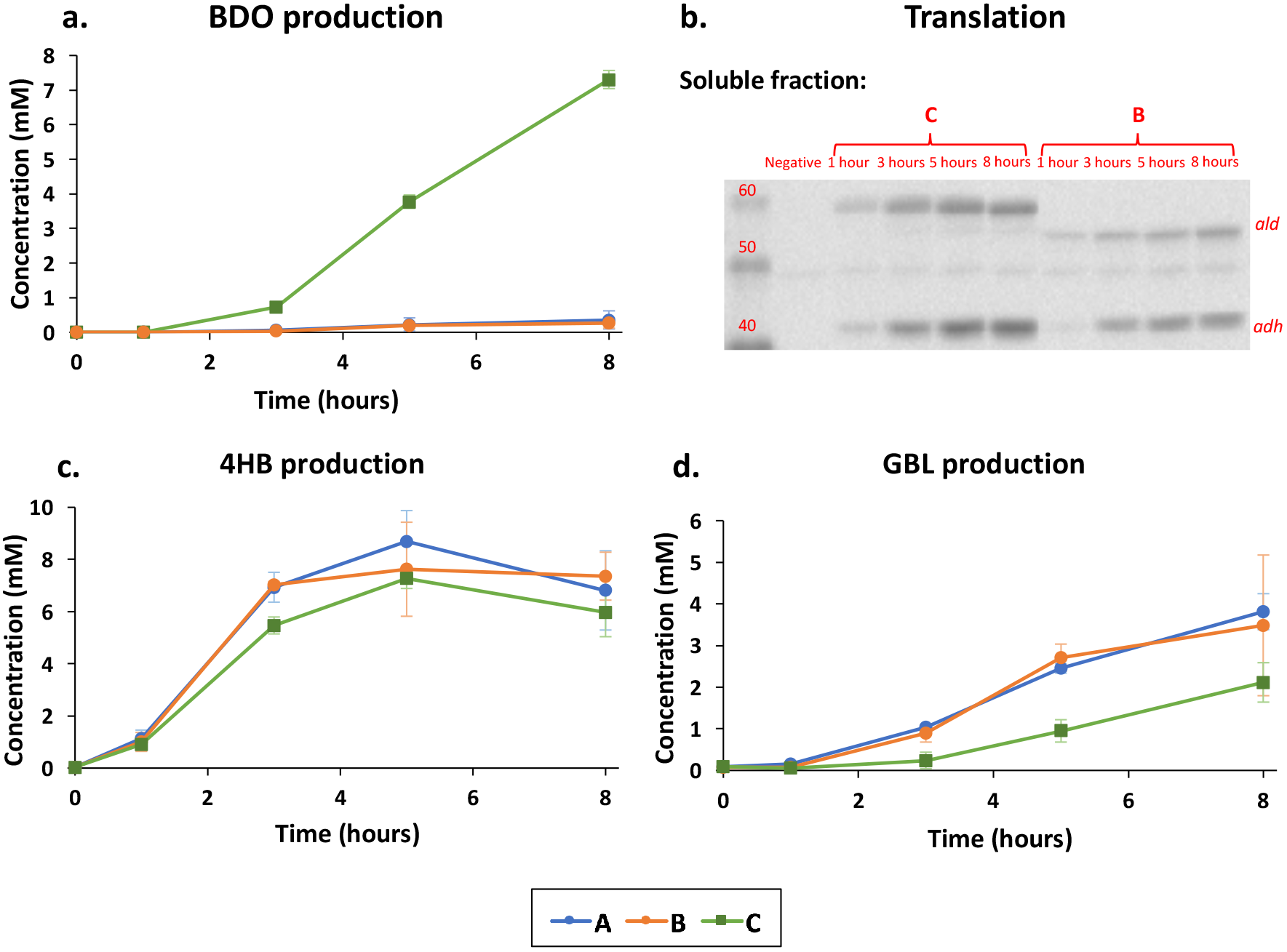
Application of TX-TL—pathway ranking. a. BDO production in the first 8 hours of the TX-TL reaction is shown. b. Soluble protein in the first 8 hours of the TX-TL reaction is shown. c. 4HB production in the first 8 hours of the TX-TL reaction is shown. d. GBL production in the first 8 hours of the TX-TL reaction is shown. Blue circle—enzyme combination A, orange circle—enzyme combination B, green square—enzyme combination C. Error bars show one standard deviation for n ≥ 3 independent experiments.

### Conclusion

This work shows that TX-TL can be a reliable engineering platform for metabolic engineering. Through studying transcription, translation, and metabolism in TX-TL, we find that the limitation of the TX-TL system lies within rapid degradation of mRNA and cofactor imbalance. The main energy source of TX-TL is glutamate, which is different from the *in vivo* system but this does not impact its ability to produce target metabolites similar to small-scale *in vivo* synthesis. The ease of adding DNA of interests and tuning protein expression by DNA concentrations opens up unique opportunities for rapidly exploring pathway design space. Studying the BDO pathway in both TX-TL and *in vivo* helps confirm that TX-TL is a reliable tool for capturing pathway dynamics. Successfully demonstrating the viability of transcription and translation machinery in S138 extract and the production of BDO provides confidence for future prototyping and engineering in extracts made with advanced *E. coli* strain or potentially other organisms.

## Materials and Methods

### Cultivation

48-well plates are used for cell culture and metabolite production. M9 minimal salts medium (6.78 g l^−1^ Na_2_HPO_4_, 3.0 g l^−1^ KH_2_PO_4_, 0.5 g l^−1^ NaCl, 1.0 g l^−1^ NH_4_Cl, 1 mM MgSO_4_, 0.1 mM CaCl_2_) is used, and it is supplemented with 10 mM NaHCO_3_, 20 g l^−1^ D-glucose and 100 mM MOPS to improve the buffering capacity, 10 µg ml^−1^ thiamine and the appropriate antibiotics. 0.5 mM of IPTG is added to induce enzyme expression. Cell culture starts with overnight growth in LB followed by another overnight growth in minimal media and final culture. The final culture is placed on a shaker at 650 rpm at 30°C or 37°C for cultivation. The culture is centrifuged to collect pellet for protein analysis and supernatant for metabolite analysis.

### Cell-Free Expression Preparation and Execution

Fermentations are performed with 1 L initial culture volume in 2-l Biostat B+ bioreactors (Sartorius Stedim Biotech). The temperature is held at 37 °C, and the pH is held at 7.0. Cells are grown aerobically to an optical density (O.D.) of approximately 10, at which point the culture goes through a series of spin-down, re-suspense, and wash cycle according to protocol described by colleagues previously.^41^ Collected cell pellets are homogenized using M-110F Microfluidizer Processor and extracted according to the method described by Kwon *et al.*^42^ with the post-homogenization incubation period extended to 80 min instead of 60 min. Buffer preparation is done according to Sun *et al.*’s protocol^41^ with a supplement of 30 mM maltodextrin. The preparation results in extract with conditions: 8.9–9.9 mg/mL protein, 4.5–10.5 mM Mgglutamate, 20-40 mM K-glutamate, 0.33–3.33 mM DTT, 1.5 mM each amino acid except leucine, 1.25 mM leucine, 50 mM HEPES, 1.5 mM ATP and GTP, 0.9 mM CTP and UTP, 0.2 mg/mL tRNA, 0.26 mM CoA, 0.33 mM NAD+, 0.75 mM cAMP, 0.068 mM folinic acid, 1 mM spermidine, 30 mM 3-PGA, 4 mM NADPH, 1 mM acetyl-coA, and 1 mM NADH. When needed, inducers such as IPTG, linear DNA or plasmid DNA are added to a mix of extract and buffer. TX-TL reactions are conducted in PCR tubes and kept at 29°C with incubation in PCR machine. BioTeK Synergy H1 microplate reader is used to collect kinetic data for fluorescent protein and MG-Aptamer.

### Plasmid DNA, PCR Product Preparation, and Cloning

PCR products are amplified using KOD (Novagen, EMD). Plasmids are miniprepped using a QIAprep spin columns (Qiagen) and further concentrated with Millipore’s Amicon Ultra-0.5 Centrifugal Filter Unit with Ultracel-30 membrane. All plasmids are processed at stationery phase. Before use in the cell-free reaction, PCR products undergo an additional PCR purification step using QIAquick PCR Purification Kit (Qiagen), which removes excess salt detrimental to TX-TL and are eluted and stored in water at −20°C for long-term storage. Gibson Assembly Ultra and Gibson Assembly HiFi Master Mix from SGI-DNA are used for plasmid assembly.

### Analytical Methods

LCMS/MS is used to analyze TX-TL samples and supernatant of cell culture. TX-TL samples were first diluted with 1:3 volume ratio with methanol to remove proteins and other big molecules, and then diluted 1: 12.5 with diluent containing labelled internal standards before LCMS analysis. Cell culture samples were diluted 1:50 with same diluent before LCMS analysis. API3200 triple quadrupole system (AB Sciex, Life Technologies, Carlsbad, CA), interfaced with Agilent 1260 HPLC, utilizing electrospray ionization and MRM based acquisition methods is used. BDO and GBL are detected using positive ionization mode, while 4HB and related acidic compounds are detected using negative ionization mode. Zorbax Eclipse XDB C18 4.6x30mm (particle size 1.8um) was used. Column temperature is maintained at 40°C, flow rate of 0.7 mL/min. Injection volume is 5 ul. Eluents include water with 0.1% formic acid (A) and methanol with 0.1% formic acid (B). A fast 1.5 min 5-95% methanol gradient is used, resulting in 1.5 min long LCMS method.

### Protein Gel and Western blotting

Culture pellets are resuspended with Novagen BugBuster Protein Extraction solution mixed with rLysozyme and benzonase. The amount of BugBuster added is normalized based on the OD value of 1mL culture. Samples are incubated at room temperature for 20 minutes. An equal volume of 2X Laemlii buffer containing 5% of ß-mercaptoethanol is mixed with each sample. The mixture is subsequently boiled in a thermocycler at 99°C for 5 min. 10ul of a mixture containing cell pellets is loaded into each lane, or 5 ul of a mixture containing TX-TL sample is loaded. Invitrogen MagicMarker XP Western Protein Standard and Bio-Rad Kaleidoscope is added as protein ladders. Bio-Rad 4-15% Criterion TGX (Tris-Glycine eXtended) Stain-Free precast gels are used. Gel imaging is done using Bio-Rad Gel Doc EZ Imager for measuring total protein intensity. Invitrogen iblot system is used for gel transfer. Western blot imaging is done using the Chemi Hi Resolution setting on a BioRad ChemiDoc MP imager. The intensity from Western blot is normalized by the total protein from each lane, and stain-free total protein is the loading control for Western Blots.^37^ Protein band intensity is determined by using Image Lab 5.2.1.

## Acknowledgement

We thank Dr. Nathan Dalleska and the Environmental Analysis Center for the support and assistance using GC/MS for initial data collection. We thank Anna Lechner, Evan Ehrich, Emily Mitchell, Glenn Majer, and Sari Rizek for material analysis and preparation. We thank Jingyi Li, Jungik Choi, Jonathan Joaquin, Joseph R. Warner, Robin Osterhout, and Robert Haselbeck for helpful discussion. This material is based upon work supported in part by the Defense Advanced Research Projects Agency (DARPA/MTO) Living Foundries program; contract number HR0011-12-C-0065 (DARPA/CMO). Y.Y.W was supported by NIH/NRSA Training Grant 5 T32 GM07616 and the Gordon and Betty Moore Foundation Grant GBMF2809 to the Caltech Programmable Molecular Technology Initiative.

